# Evaluating the effectiveness of raised beds in alleviating heavy metal contaminants

**DOI:** 10.64898/2026.11.13.766404

**Authors:** Xorla S. Ocloo, Nicholas Medina

## Abstract

Structural policy and social practices, including redlining and white flight, have driven disinvestment and heightened exposure to persistent environmental pollutants resulting from residual impacts of industrial activity. Urban agriculture, meant to empower and feed communities, are frequently established on vacant lots that may contain these contaminants and pose a threat to human health. To circumvent the legacy of contaminations on bare soil, Chicago growers are using raised beds to create a barrier between contaminated soil and healthier imported compost. Our goal was to assess the efficacy of this barrier system by 1) quantifying eight heavy metals and Polychlorinated Biphenyls (PCBs) between existing soils and raised beds, 2) examining the difference between key soil properties, and 3) determining the environmental risk determined by their potential ecological risk index (PERI). We expected that metal concentrations would vary across sites and soil types, and raised beds would have fewer heavy metals than existing soils. Our results showed existing soils generally had higher levels of heavy metals with significant differences in lead and arsenic, suggesting that raised beds could be a viable barrier system. We also found higher levels of phosphorus, nitrogen, and magnesium in raised beds than existing soils. Thirdly, lead and copper significantly increased with pH while arsenic and copper decreased with organic matter levels. Despite the presence of heavy metals, the sites had a lower risk of environmental contamination, indicating that growers’ extra measures to ensure their soils are safe to grow food may be sufficient, barring metal- and site-specific strategies.

**PLAIN LANGUAGE:** Urban areas used as farms or community gardens are susceptible to toxic substances such as heavy metals and Polychlorinated Biphenyls (PCBs) due to industrial activity. However, there are ways to mitigate exposure by using raised beds. In this study, we analyzed 8 different types of heavy metals, PCBs, and key soil properties such as pH and organic matter to determine differences between existing soils and bed soils. We found that existing soils had higher levels of lead and arsenic compared to soils found in raised beds. We also found lead and copper significantly increased with pH while arsenic and copper decreased with organic matter levels. This suggests that using raised beds could be a safe and effective strategy when growing on land with a history of soil contamination.

## INTRODUCTION

Urban agriculture (UA) has resurged in countries in the Global North in hopes of reviving neglected neighborhoods (Lovell & Johnston, 2009) by increasing employment, providing economic empowerment, and addressing food apartheids through food sovereignty (Gripper et al., 2022; Mkwambisi et al., 2011). UA not only provides more equitably distributed and healthy foods, but a place to gather for social and cultural functions that strengthen community ties (Lovell, 2010). Moreover, UA provides a plethora of ecological benefits for urban core areas such as increase in biodiversity, air, soil, and water quality (Orsini et al., 2013). Despite great social and ecological benefits often found in urban agriculture, urban soils experience environmental pollution due to long industrial legacies of contaminated waste in cities. Consequently, this directly threatens public health and food safety for exposed local growers and community members (Oka et al., 2014).

Oftentimes, UA sites are found on vacant lots that are contaminated with metals and chemical products deposited by previous industrial activity (Oka et al., 2014) and ongoing automobile traffic emissions (Kim et al., 2014). This legacy pollution, most often found in Black and Brown communities, are then converted into community gardens where growers must remediate the land before using (Grifoni et al., 2022). For example, Cadmium (Cd), Molybdenum (Mo), Nickel (Ni), Zinc (Zn), Arsenic (As), and Lead (Pb) are notable metals from sources of commercial fertilizers, liming materials, agrochemicals, sewage sludge and irrigation water (He et al., 2005). Most at risk are growers and consumers via direct contact, ingestion, or inhalation of contaminated soil particles and crop plants (Dudka et al., 1996; McBride et al., 2014; Spliethoff et al., 2016). Additional risk accumulates when industry escapes remediation responsibility and the burden of expensive strategies, such as replacing topsoil or soil washing (Noller et al., 2022), falls on oppressed communities. Therefore, it is important to assess heavy metal contamination and ongoing risks in growing community gardens and urban farms across cities, including how they vary within cities.

Globally, Chicago is one of the most populated, yet segregated and polluted cities, where the legacy of redlining and industrial pollution continues to shape the current landscape of environmental health (Kumar & Hundal, 2016). Specifically, Pb has persistently been and continues to be a growing concern in urban sites, maintaining an alarming health risk for urban agriculture (Brown et al., 2016; Cheng et al., 2015). Watson et al. (2022) found that concentrations of Pb ranged from 10 parts per million (ppm) to 1600 ppm and with samples averaging at 220 ppm across the city of Chicago, an average which exceeds federal US EPA’s 200mg/kg limit. Another investigation found that more than 54% of the soil samples collected exceeded this EPA limit, with Pb hotspots in the south and west sides of Chicago (Thorstenson et al., 2025). However, despite this widespread contamination and drawing from the need to combat food apartheid, there are over 800 self-reported growing sites throughout Chicago, most of which are sitting on land that were previously exposed to contaminants (Growing Food and Open Space in Chicago, 2022). Furthermore, polychlorinated biphenyls (PCBs) were widely used in industrial and manufacturing companies in electrical transformers, capacitors, caulk and joint sealers, electrical wiring coatings, carbonless copy paper, and adhesives between 1929 and 1979 (Erickson & Kaley, 2011; Shanahan et al., 2015). They are environmental pollutants that are resistant to chemical, thermal, and biological degradation (Shanahan et al., 2015). Soils may be the largest global PCB repositories due to deposition from manufacturing, leaching from building materials or landfills, and the application of wastewater treatment plant biosolids (Meijer et al., 2003). With newer community gardens emerging in the south and west sides of Chicago, grower efforts are threatened by the risks of persistent exposure to heavy metal and PCB contamination. Establishing these growing sites often requires costly remediation that can still leave behind lingering threats to ecosystems and communities (Paltseva et al., 2022).

Soil pH, compost, and nutrients play a role in heavy metal distribution in urban soils (Li et al., 2022; Medyńska-Juraszek et al., 2020; Young, 2013). When soil becomes more acidic, heavy metals are more mobile and easily available for plant uptake (Gruszka et al., 2025). A neutral pH can help keep heavy metal concentrations low (Cheng et al., 2011). Compost, which is used in raised beds, typically has higher levels of nutrients such as Phosphorus (P), Nitrogen (N), and Potassium (K) and other organic matter than most native topsoils (Feng et al., 2021). Though high in macronutrients, this can also increase Cd concentrations while decreasing Pb concentrations (Clarke et al., 2015). Alternatively, soil organic matter, which plays a critical role in key soil functions chemical processes, can bind metals into insoluble forms, limiting their movement (Fidos & Rutkowska, 2023).

To address potential risks of soil contamination, growers are using alternative growing methods to ensure safe food production. Urban agricultural sites commonly use organic practices such as adding compost for remediation (Attanayake et al., 2014, 2017; Defoe et al., 2014) and companion cropping. Moreover, there has been a widespread interest in building raised beds, instead of direct row sowing, which brings in high-quality soils, limits exposures to soil contaminants, and has indigenous origins (Pohl et al., 1996; Sullivan et al., 2015). Typically, a raised bed media includes a mixture of soil, compost, perlite, and/or sand which helps retain water for plants (Agnew & Leonard, 2003). Raised beds also improve conditions for water drainage (Takeshima et al., 2023) and generally have higher nutrient content (Lopez-Mondejar et al., 2010). Nelson et al., (2022) found trends toward higher active carbon, C/N, and sulfur in Oregon raised beds than in-ground garden beds, though not significant. Permanent raised beds also improved soil structure (i.e., macro-aggregate stability, aeration and capillary porosity, and soil moisture) and crop yields compared with traditional tillage (Araya et al., 2020). A study in Chicago found that raised beds showed lower levels of Pb than bare areas, suggesting that raised beds help alleviate contamination for other heavy metals (Witzling et al., 2010). While much research quantifies heavy metals in Chicago (Jenkins et al., 2026; Watson et al., 2022), less is done on investigating the heavy metals and important soil health factors between barriers, such as raised beds which are meant to separate growing beds from potential contaminants in existing soils. Specifically, nuances may exist between certain heavy metals, such as Pb which is more responsive than As (McBride et al., 2014). Additionally, heavy metals in dust can lead to recontamination even after amendment in the field, as shown in Boston (Clark et al., 2008), which may undermine otherwise beneficial effects of UA amendments (Miernicki et al., 2018).

In this study, we investigated a wider array of heavy metal contamination and associated them with soil chemical fertility levels across seven Chicago community gardens and urban farms. Our objectives were to 1) quantify eight heavy metals and Polychlorinated Biphenyls (PCBs) between existing soils and raised beds, 2) examine the difference between key soil properties in the two soil types, and 3) determine the overall environmental risk at each of the seven sites as determined by their potential ecological risk index (PERI). Considering Chicago’s industrial past, we expected heavy metal and PCB contamination at all sites, but that barriers, such as raised beds, would help mitigate potential health risks. We expected that raised beds across all sites will have higher soil nutrient concentrations than existing soils. We also expected that heavy metals may be associated with soil nutrient levels, with organic matter alleviating heavy metals (Cao et al., 2003). Overall identifying contamination in Chicago community gardens and urban farms is important in highlighting a promising approach that can used in urban agricultural systems worldwide that is safe and productive despite a legacy of soil contamination.

## METHODS

### Study sites

We studied seven urban agricultural sites in Chicago’s southside: four community gardens and three urban farms. The precise locations and names of the sites are not identified to maintain confidentiality. We selected sites based on developed community partner relationships, active and responsive stewardship, focusing on neighborhoods with high proportions of diverse people and lower median incomes. Most of the raised beds across sites were filled using the Hügelkultur layering method which included thick branches at the bottom, green wastes, and compost as the top layer, similar to other strategies (Moskal & Berthrong, 2018). During collection, farmers were currently growing flowers (i.e., sunflowers and marigolds), herbs, and vegetables (e.g., kale, peppers, carrots, scallions, brussels sprouts, eggplant, etc.). Many sites were adjacent to roads, churches, food pantries, housing, restaurants, and near train tracks.

#### Sample Collection

Soil samples were collected in November and December of 2024, when daily temperature varied between 0-8°C with some snowfall events. Sampling targeted raised beds and adjacent existing soils in areas where managers deemed important based on their concerns or reservations for growing. Garden C did not have any exposed existing soil. Thus, we only collected samples from the raised beds. Garden G was the only location that also had soil that were not remediate. Therefore, we collected some samples to evaluate the differences between non-remediated and remediated existing soils. We collected the top 10cm of soil which is the main rooting depth for most vegetables (Chang et al., 2014) with a hand shovel that was washed between samplings. For each sample, approximately 300g of soil was collected and composited into Ziplock bags (Malone et al., 2023). Thirty-four total samples were collected and composited by site and soil source (i.e. raised bed or existing).

#### Soil chemical analysis

In total, twenty-three composite samples were analyzed at the Pennsylvania State University Agricultural Analytical Service Lab. We tested the samples for Arsenic (As) EPA Method 3050B + 6010, Cadmium (Cd) EPA Method 3050B + 6010, Copper (Cu) EPA Method 3050B + 6010, Molybdenum (Mo) EPA, Nickel (Ni), Zinc (Zn), and Lead (Pb) (EPA Method 3050B + 6010), as well as Mercury (Hg) (EPA Method 7473) and Molybdenum (Mo) EPA Method 3050B+ 6010, Nickel (Ni) EPA Method 3050B + 6010, Zinc (Zn) EPA Method 3050B + 6010, polychlorinated biphenyls (PCBs) (EPA Method 8082). Samples were also analyzed for pH, total nitrogen (combustion), cation exchange capacity (CEC), extractable Mehlich 3 P, K, Ca, and Mg, as well as and organic matter using loss on ignition at 55°C method SM 2540 G. (Compost Methods Agricultural Analytical Services Lab, 2026).

#### Statistics

All analyses were done in R v4.4 (R Core Team, 2022). We analyzed the average concentration and SE of all metals across the seven sites. We ran a spearman correlation analysis to observe the relationship between all metals and fertility variables, and none were strongly correlated above 0.4 and thus treated independently in downstream analyses. For soil fertility and metal concentrations, we compared differences between existing and bed soil sources in each relevant variable using non-parametric wilcoxon tests with the *wilcox_test()* and *wilcox_effsize()* functions in the *rstatix* package, and tested site effects using *kruskal_test()* and *kruskal_effsize()* in the same package. This approach is used in similar studies (Edwards et al., 2023) and resembled analogous generalized linear models. Linear regressions between soil nutrient and heavy metal data excluded clear outliers and were run using *stat_smooth()*.

Additionally, Hakanson (1980) proposed a method of evaluating heavy metal pollution in sediments, which comprehensively considers the toxicology and the ecological and environmental effects of heavy metals. It was later widely used in the evaluation of heavy metal population in soils (Zang et al., 2017). The equation is as follows:

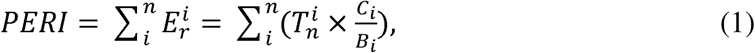

where C*_i_* and B*_i_* are the measurement concentration and local background value, respectively, of soil heavy metal *i* [25], Ti n is the eco-toxicity response coefficient of heavy metal *i*, and E*i r* is the PERI of heavy metal *i*. We used this equation to calculate the ecological risk of heavy metals at all sites. The potential ecological risk index can be classified into four categories: low (PERI < 150), moderate (150 ≤ PERI < 300), high (300 ≤ PERI < 600) and very high (PERI ≥ 600) (Huang et al., 2021).

## RESULTS

### Soil fertility

We observed relatively high levels of soil fertility across all urban agriculture sites, and significant differences between existing and raised bed soils across all measured variables (*P* < 0.05) except soil organic matter and marginal significance in pH (Table 1). Broadly, each source of soil tended to differ most (*r* > 0.7) in P (*W* = 0.5, *P* < 0.001) and Ca (*W* = 10, *P* < 0.001); the least (*r* < 0.5) in pH (*W* = 96, *P* = 0.0584) and K (*W* = 30, *P* < 0.05), and substantially in all else including N (*W* = 11.5, *P* = 0.001), CEC (*W* = 11, *P* < 0.001), and Mg (*W* = 12, *P* < 0.001). Specifically, bed soils were 0.15 units more acidic than existing soils. Existing soil organic matter was relatively high around 7%, and bed soils contained twice that amount. Bed soils also contained twice as much N, Mg, and nearly five times as much P. Trends showed notable, but slightly smaller differences in CEC and Ca of about 20% higher in raised bed soils. Despite minor variation (Table 1), we did not detect significant effects of garden site on soil fertility metrics.

**Table 1.** Measures of fertility between native urban and raised bed soils across various urban agriculture sites (Chicago, IL, USA); including pH, soil organic matter and nitrogen (N) in percent units, cation exchange capacity (CEC) in centimoles per kilogram, and phosphorus (P), calcium (Ca), and magnesium (Mg) in units of parts per million (ppm).

| Site Label | Soil Source | pH | OM pct | N pct | P ppm | CEC | K ppm | Mg ppm | Ca ppm |
|---|---|---|---|---|---|---|---|---|---|
| Garden_A | Existing | 7.0 | 16.7 | 0.9 | 163.0 | 20.7 | 206.0 | 622.0 | 4,478.3 |
| Garden_A | Raised Bed | 7.3 | 10.1 | 0.5 | 248.0 | 22.3 | 260.0 | 791.0 | 4,656.3 |
| Garden_B | Existing | 7.4 | 15.4 | 0.7 | 125.0 | 21.3 | 185.0 | 701.0 | 4,061.1 |
| Garden_B | Raised Bed | 7.2 | 13.1 | 0.9 | 366.7 | 23.0 | 390.3 | 837.0 | 6,696.2 |
| Garden_C | Raised Bed | 7.2 | 14.7 | 1.1 | 686.5 | 25.2 | 612.0 | 1,032.5 | 5,194.8 |
| Garden_D | Existing | 7.5 | 12.2 | 0.4 | 109.5 | 21.0 | 287.5 | 634.5 | 4,587.6 |
| Garden_D | Raised Bed | 7.3 | 16.1 | 1.3 | 475.7 | 25.2 | 622.0 | 1,028.7 | 5,124.8 |
| Garden_E | Existing | 7.5 | 21.4 | 0.8 | 107.0 | 22.0 | 478.0 | 691.0 | 4,036.1 |
| Garden_E | Raised Bed | 7.5 | 13.4 | 0.8 | 262.5 | 22.4 | 417.5 | 754.5 | 4,478.2 |
| Garden_F | Existing | 7.7 | 6.8 | 0.3 | 143.0 | 19.9 | 267.0 | 506.0 | 4,284.4 |
| Garden_F | Raised Bed | 7.3 | 14.8 | 0.9 | 518.0 | 21.4 | 275.0 | 684.0 | 5,130.1 |
| Garden_G | Existing | 7.6 | 4.5 | 0.3 | 54.8 | 17.9 | 191.8 | 390.5 | 3,645.4 |
| Garden_G | Raised Bed | 7.5 | 24.8 | 1.4 | 571.0 | 28.7 | 1,433.0 | 1,199.0 | 4,825.0 |

### Heavy metals and PCBs

We analyzed all eight metal concentrations across existing vs. raised bed soil types and found heterogeneity across our seven sites (Fig. 1). None of the sites exceeded the Cook County background soil limits for As. Three of the sites exceeded the background levels for Ni. Five out of the seven sites exceeded the background limits for Cd and Hg. All sites exceeded the background levels for Cu and Zn. Only one site exceeded the US EPA limits for Pb of 200 ppm (US EPA, 2024). Overall, existing soils tended to show higher levels of all metals, with significant differences compared to bed soils in Pb (*W* = 100, *P* = 0.03), As (*W* = 99, *P* = 0.036), and both available Cu (*W* = 107, *P* = 0.008) and available S (*W* = 30, *P* = 0.03), with no detectable differences between soil sources in other metals. Importantly, focusing on Pb, existing soils at one location tended to exceed the US EPA threshold safety levels, reaching up to 600 ppm, while bed soils hovered at approximately 100 ppm, just above the California standard of 80 ppm (State of California, 2020).

**Fig. 1.**
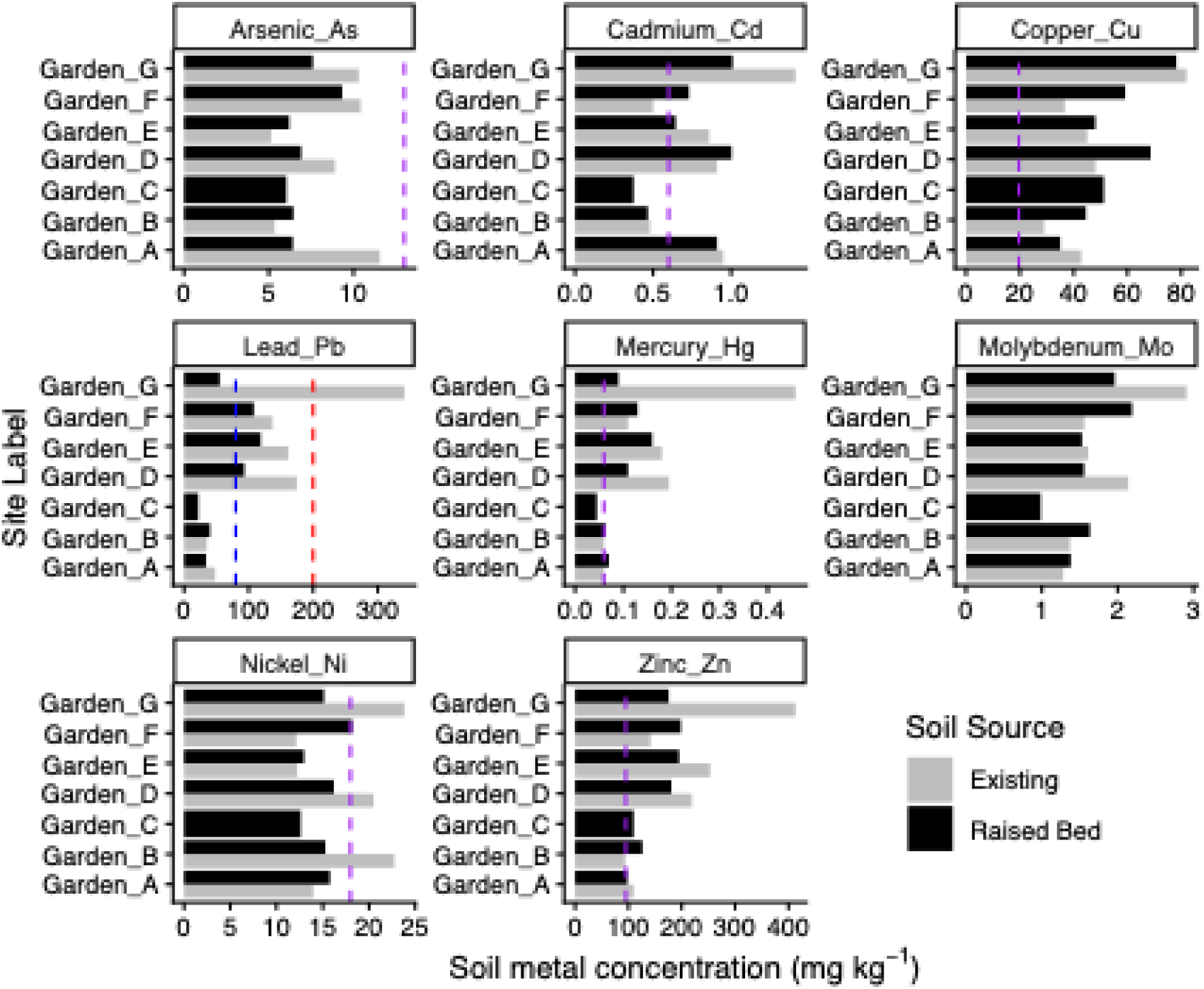
Heavy metal concentrations, in units of mg per kg^-1^, between existing (light grey) and raised bed (dark grey) soils (IL, USA), including Arsenic, Cadmium, Copper, Lead, Mercury, Molybdenum, Nickel, and Zinc. The red dotted line represents the EPA limits, and the blue dotted line represents the California limits, which matches UK limits (Entwistle et al., 2019). The purple dotted line represents Illinois metropolitan background soil limits based on the Tiered Approach to Corrective Action Objectives (TACO), codified under 35 Ill. Adm. Code 742, and set by Illinois Environmental Protection Agency (Illinois Environmental Protection Agency, 1997). Molybdenum did not have soil background limits in metropolitan Illinois.

We observed marginally significant effects of UA site on Pb (*X*^2^ = 12.1, *P* = 0.06), As (*X*^2^ = 12.5, *P* = 0.0521), Hg (*X*^2^ = 12.1, *P* = 0.0598), Cd (*X*^2^ = 12.2, *P* = 0.056), and Zn (*X*^2^ = 10.7, *P* = 0.098) (Fig. 1). In general, we found that Garden C had the lowest concentration of all eight metals and Garden G had the highest concentration of all metals. Four out of seven sites exceeded the California safety levels of Pb in the soil, while only one location, the existing soil at Garden G, exceeded the EPA safety levels for Pb at 341.09 mg/kg^-1^). For As, all growing sites had at least 6 mg/kg^-1^ in their soil types. Cd concentrations were highest at Garden D and G with over 1 mg/kg^-1^). We found the lowest concentration levels of copper at Garden B, A, and F, all averaging under 37 mg/kg^-1^. For Hg, we found that Garden A, B, and C had relatively low levels, whereas Garden D and G had higher levels. Garden C had the lowest Mo concentration at 0.99 mg//kg^-1^, whereas the highest were found in the existing soil at Garden G and Garden D (over 2mg/kg^-1^). We found the lowest concentration of Ni in the raised bed soil and existing soil at Garden E and in the existing soil at Garden C. We found that the raised bed soil at Garden A and existing soil at Garden B had a Zn concentration level lower than 100 mg/kg^-1^. Garden G had the highest level of Zn found in their existing soil (413.07 mg/kg^-1^). Lastly, six out of seven sites had PCB concentrations of less than 0.02 mg/kg^-1^. The existing soil at Garden G was the only location where PCB concentrations were 0.07 mg/kg^-1^ on average.

### Soil metal correlations

Soil metals were significantly predictable from soil fertility properties, namely either soil pH and/or soil organic matter (Fig. 2). Specifically, higher pH was associated with more Pb and Cu, where increasing pH by 0.1 increased Pb by 22 ppm (*p* = 0.001, *R^2^* = 0.44) and Cu by about one (*p* = 0.0006, *R^2^* = 0.33). In contrast, soils with more organic matter lowered soil As (*p* = 0.013, *R*^2^ = 0.26) and Cu (*p* = 0.023, *R*^2^ = 0.24) each by about one ppm per 5% increase in soil organic matter.

**Fig. 2.**
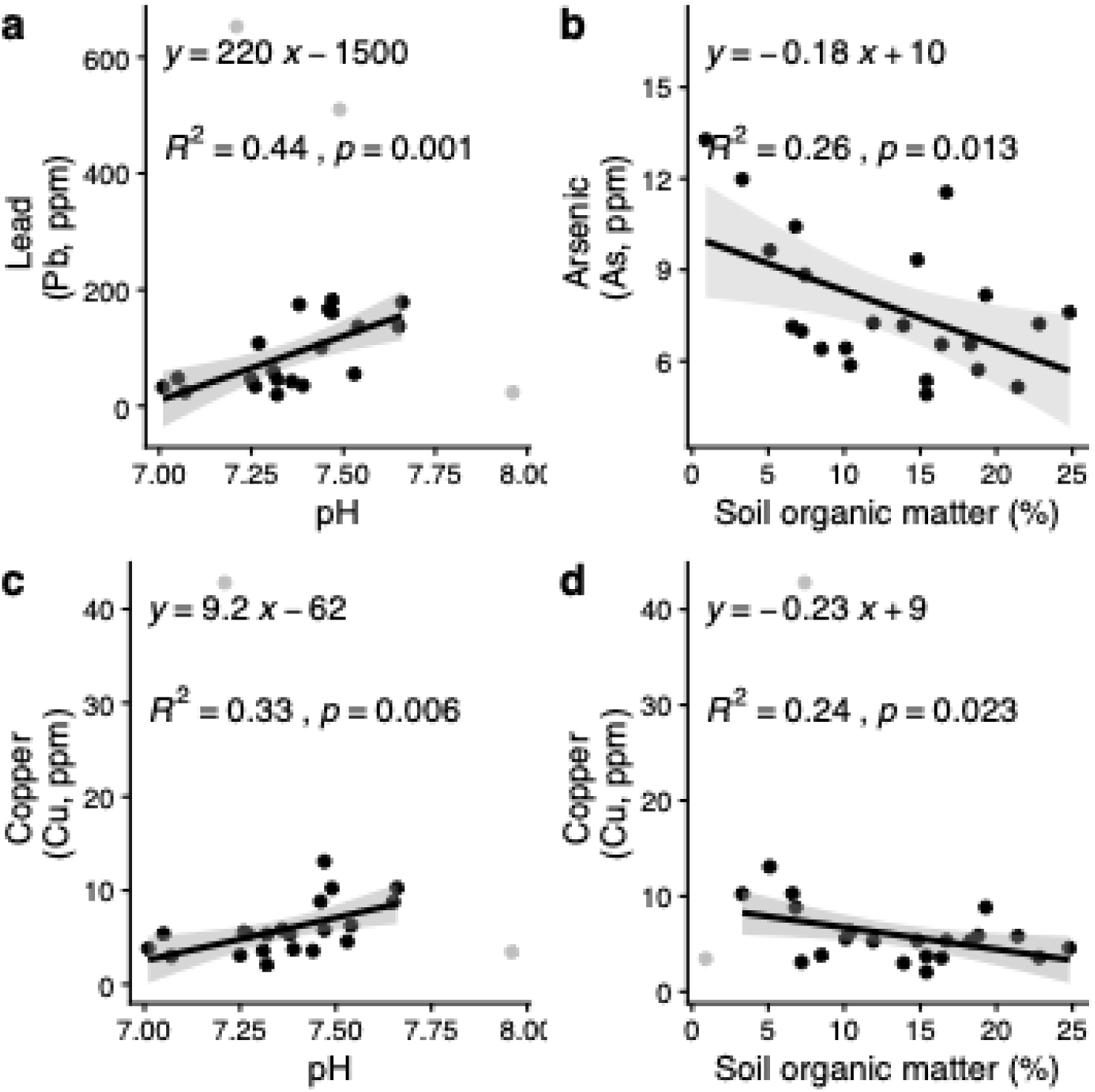
Significant soil metal (a) Pb, (b) As, and (c,d) Cu correlations with pH and soil organic matter across raised bed and existing soils from seven urban gardens and farm in Chicago, USA. Raw data (black) and excluded outliers (grey) are shown.

### Potential Ecological Risk Index

We found the highest potential ecological risk index (PERI) at Garden G (value = 306.41), located furthest South, and the lowest PERI at Garden C (value = 37.08). Gardens A and B also had a low ecological risk, values = 117 and 85, respectively. All other locations (D, E, F) had a moderate ecological risk of heavy metals, values between 153 to 196 (Fig. 3).

**Fig. 3.**
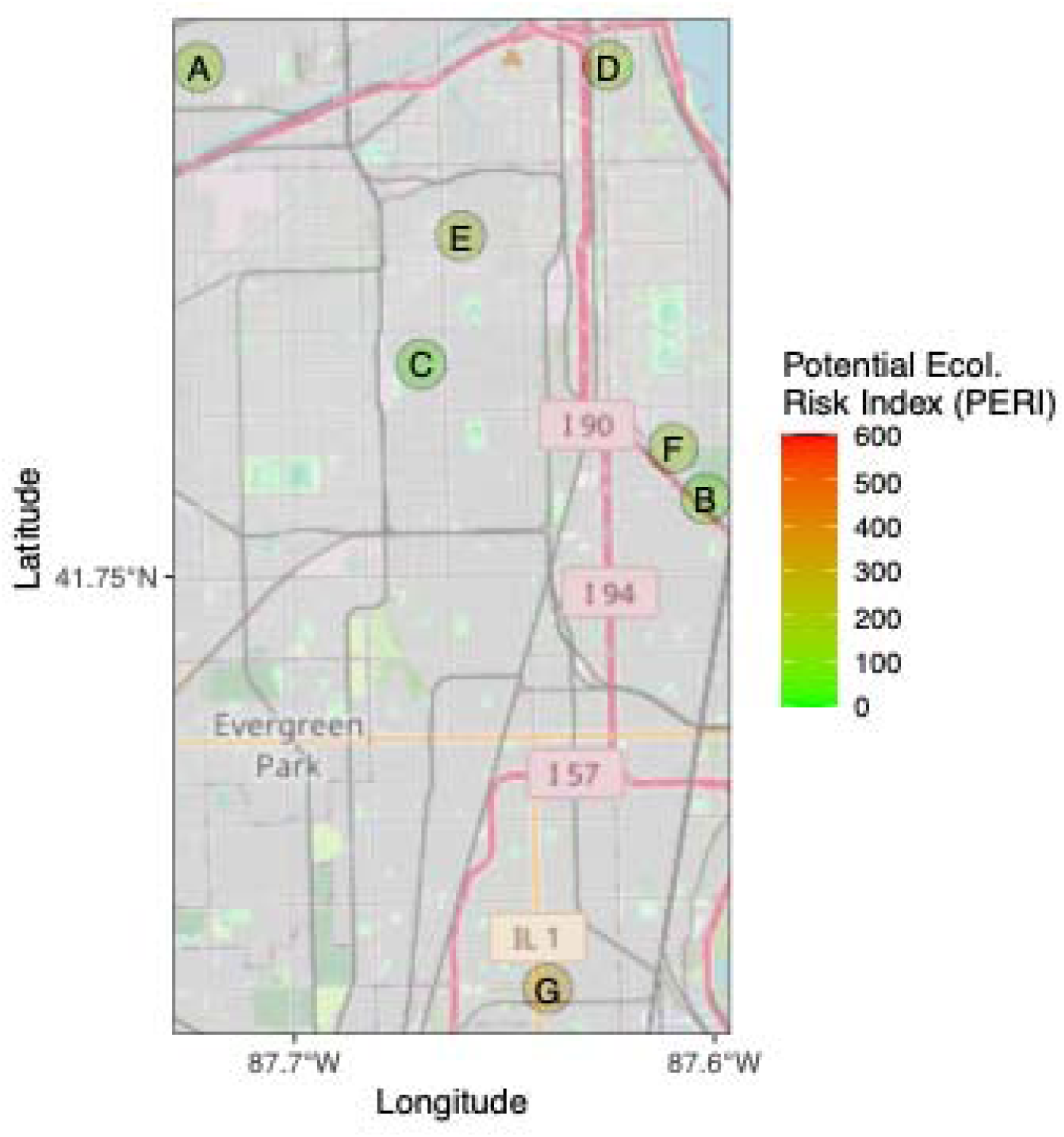
Heat map representing the various PERI across all garden sites. The different PERI rating levels are as follows: PERI < 150 = Low comprehensive potential ecological risk; 150 ≤ PERI < 300 = Moderate comprehensive potential ecological risk; 300 ≤ PERI < 600 = high comprehensive potential ecological risk; PERI > 600= very high comprehensive potential risk.

## DISCUSSION

The objective of this study was to examine how raised beds help mitigate heavy metal contamination and improve key soil properties between existing soils and raised beds at seven urban agricultural sites in Chicago. Despite limited sample size due to budget constraints, this study finds significant variation in urban garden soil metals and chemical fertility, including between existing and garden bed soils, highlighting context-dependent effects of using raised beds to alleviate heavy metal burdens. In partial support of our initial hypothesis, raised bed soils helped alleviate Pb and As concentration levels, as well as available Cu and available S, but not other heavy metals. This is in support to another study that found that barrier strategies such as mounding or using raised beds appear to be effective at reducing As and Pb in growing beds (Moskal & Berthrong, 2018). We also detected spatial variation in five different metals, despite similar soil fertility across UA sites.

Nutrients in plants are essential for proper growth and development. Most importantly, plants require high amounts of macronutrients such as nitrogen (N), phosphorus (P), and potassium (K) which play a role in important metabolic processes and aids photosynthesis (Kulcheski et al., 2015). At four of the six sites (Gardens A, B, D-G), N was highest in the raised beds. All sites excluding Garden C, had a higher P level in the raised beds, similar to UA in NYC (Mitchell et al., 2014). Lastly, raised beds showed the highest level of K at five of the six sites. Out of these three macronutrients, P was significantly higher in raised bed soils than existing soils. This finding indeed aligns with those of similar previous studies, showing excess P concentrations beyond 80-100 ppm accumulating in existing soils under garden beds as well (Small et al., 2019) up to at least seven years (Small et al., 2024). However, soil P did not predict any heavy metal concentration across sties, perhaps due to consistency in high P across sites (Cao et al., 2003; Obrycki et al., 2016). Similar soil fertility across sites may reflect regional sources of used raised bed soils, as well as similarly depleted urban soils facing urban heat island and limited moisture effects (Shi et al., 2012). Moreover, the optimal phosphorus level for plant and crop production generally ranges between 30 to 50 parts per million (ppm) in soil tests (Penn State Agricultural Analytical Services Lab, 2026). All sources of soils exceeded the optimal phosphorus level suggesting that agricultural sites have excess P that could leach into nearby waterbodies and cause eutrophication. Although, certain organic practices introduce important macronutrients, too much can also raise environmental issues. Raised beds also showed higher calcium and magnesium than existing soils underneath. This could be because most growers are using soil amendments that are usually higher than native soils to promote plant growth (Penn State Agricultural Analytical Services Lab, 2026).

The heterogeneity in heavy metals across UA sites can also be explained by several factors. Broadly, higher metal concentrations in existing soils compared to raised beds supports raised bed strategies to lower metal exposure, including Pb, but not As (Paltseva et al., 2018; Peryea, 1991; Ramazanova et al., 2026b; Scheckel et al., 2013), but may not lower them down below Illinois background soil levels, except in the case for Ni. Thorstenson et al. (2025) sampled 1,750 soil samples in Chicago and found the median soil Pb concentration to be 217 ppm. Garden G’s non-remediated soil was measured greater than 300 ppm and less than 100 ppm in the raised beds, signaling a strong support of raised beds reducing the amount of Pb. Similarly, Witzling et al. (2010) found that raised beds reduced Pb levels compared to adjacent areas of bare soil in Chicago community gardens. In contrast, despite significant variation across UA sites, raised beds may not be enough to completely alleviate the legacy Hg, Cd, Cu, or Zn concerns. For example, raised beds were effective in lowering Pb and As, but not Zn, Cu, V, and Ni in raised beds in Philadelphia community gardens (Bassetti et al., 2023). Additional strategies may include building up soils with further amendments such as compost, manures, and/or biosolids to immobilize Pb, potentially alongside sulfur to immobilize As (Paltseva et al., 2020). Emerging strategies may also include phytoremediation with metal hyperaccumulating plants, likely more useful for Ni over Pb (Bhat et al., 2025; Reeves et al., 2018), and/or myco-remediation with biomass from certain fungi, such as multi-use (Karunarathna et al., 2025; Lu et al., 2026) polypore *Ganoderma* (Parasnis et al., 2024). Interestingly, As was below the level of concern for both soil types at all sites. Ugarte and Taylor (2020) found a similar result with As, Pb, and Zn in twenty-one community garden and urban farm sites in Chicago. Arsenic may not be as high in raised beds because growers are using cedar lumber instead of pressure treated lumber which leaches chromated copper arsenate (CCA) over time (Brandstetter et al., 2000; Clarke et al., 2015).

We observed some correlations between soil chemical properties and toxic metals. Soil pH is important for vegetation growth, nutrient release, and soil microbial activity (Wang et al., 2024). Studies have shown that pH affects the mobility and solubility of toxic metal cations in soils (Niu et al., 2015; Suresh et al., 2011). Soil with low pH can facilitate the migration and availability of heavy metals (Chen et al., 2024). In contrast, we found soil pH was positivity correlated with an increase in Pb and Cu. Medyńska-Juraszek & Jadhav (2022) found that pH increased in Cu-and Pb-spiked soil when microplastics were added suggesting that the presence of microplastics increased soil pH and plays a role in the heavy metals sorption/desorption process. Further investigation is necessary to understand the role microplastics play in vectoring heavy metals and how they influence properties in soil. We also found that soils with higher organic matter decreased Cu and As concentrations. Similarly, other studies have found that higher organic matter can bind metals, reducing the heavy metal availability and subsequent uptake by plants (Cao & Wu, 2020; Zhang et al., 2018).

In general, we found low levels of polychlorinated biphenyls (PCBs) in the existing soils and raised beds despite Chicago’s legacy of PCB pollution from meat packing plants, steel and iron mills, and chemical plants. Several studies have documented elevated concentrations of airborne PCBs in Chicago (Hu et al., 2010; Sun et al., 2006, 2007). Shanahan et al. (2015) found that soils contribute 31% to PCB emissions in Chicago. Since we did not find high concentrations at our urban sites, this suggests that there was limited contact with PCBs at the site previously. The two known sites for heavy PCB contamination in Chicago is located at the Chicago Sanitary & Ship Canal (CSSC) and the Chicago Area Confined Disposal Facility (CDF) (Miller, 1998; Peverly et al., 2015). The closest site, Garden G, which had the highest average concentration of PCB at 0.07 mg/kg^-1^, is located about 12 miles from the Chicago Area Confined Disposal Facility. A closer investigation on the potential source for contamination is needed to understand other potential pathways of contamination.

Additional factors also likely shape the soil fertility and heavy metal characteristics of UA soils. For instance, the source of compost or topsoil used for raised beds, (i.e. local or commercial), can vary based on proximity and cost factors interacting with grower status. Another understudied factor can be the age of raised beds, which may allow varying times for plant-soil feedbacks (Bickford et al., 2022; van der Putten et al., 2013), including depletion of some nutrients and/or microbiome acclimation (Bryant & Bever, 2025). Related to this are management factors such as crop rotation, which helps alleviate rhizosphere and/or pest-mediated intraspecific competition via legacy effects (Robertson et al., 2024).

Most sites had a relatively low/moderate potential ecological risk index, suggesting that growers are conscious about curbing contamination within their garden soil. Most growers in Chicago are using raised beds or replacing the topsoil to grow their foods. While raised beds are not guaranteed to have safe levels of metal concentrations, they can reduce the amount of toxins compared to existing soils on the ground. Contamination of heavy metals can also come from multiple sources. For example, contamination can be airborne from nearby sources such as heavy traffic. Garden G had a higher PERI than the other sites only because the existing soils that were tested had not undergone remediation. Though it is impossible to get rid of all contaminants, growers in Chicago carefully plan out their sites by growing in barrier systems such as raised beds and actively seek soil remediation where they are going to grow food.

## CONCLUSION

Overall, this study offers insight into the heterogeneity in soil heavy metals and soil fertility across some UA sites, using Chicago as a focal large city. Specifically, we support raised beds in lowering heavy metal concentrations, especially for non-remediated sites with respect to Pb. However, additional remediation is required to reduce heavy metal exposure below Illinois state guidelines, and strategies by both growers and municipal entities should be helpful. Future studies should further parse out mechanisms of variation among UA sites operating under various ecological and socioeconomic conditions. Future directions should also consider how soil heavy metals accumulate in garden vegetables and fruits though plant roots.

## Data availability

Datasets are available from the corresponding author upon reasonable request.

## Author contributions

CRediT taxonomy

**Xorla Ocloo**: Conceptualization, Data curation, Formal analysis, Funding acquisition, Investigation, Methodology, Project administration, Resources, Supervision, Validation, Visualization, Writing – original draft, Writing – review & editing. **Nicholas Medina**: Formal analysis, Investigation, Software, Validation, Visualization, Writing – review & editing.

Conceptualization – XO; Data Curation – XO

Formal analysis – XO, NM

Funding acquisition – XO

Investigation – XO, NM

Methodology – XO

Project administration – XO

Resources – XO

Software – NM

Supervision – XO

Validation – XO, NM

Visualization – XO, NM

Writing – original draft – XO, NM

Writing – review & editing – XO, NM

## Acknowledgements

The authors would like to thank the numerous gardeners and community partners who opened their spaces to collaborative research. The authors would also like to thank the undergraduate research team: Zayn Ibrahim, Francesca Milliner, Eliza Peters, and Emily Marshall who assisted with data collection during this project. Thank you to Penn State’s Agricultural Lab for processing all samples and Dr. Christie Klimas for her comments on earlier drafts. Lastly, we would like to thank the DePaul Honors program and Environmental Science and Studies department for supporting this work.

## Declaration of Generative AI and AI-Assisted Technologies

None was used during the preparation of this work.

## Abbreviations

UA: Urban Agriculture
PERI: Potential Ecological Risk Index
PCBs: Polychlorinated Biphenyls
CCA: chromated copper arsenate

